# EEG mu rhythms and trait empathy during observation of a romantic partner’s pain with and without consoling touch

**DOI:** 10.64898/2026.09.26.754722

**Authors:** Sarah F. Love, Lauren P. Rowden, Bryten M. Ballenger, Amber L. Harris Bozer

## Abstract

**Introduction:** Observation of a partner in pain elicits empathy-related cortical responses and social support can modulate pain.

**Objective:** The aim of this study was to investigate empathy and the cortical response during observation of a romantic partner in pain with and without consoling touch.

**Methods:** A total of 40 participants (20 couples) completed validated measures of handedness, pain and empathy. Each member of the couple was randomly assigned to be either the recipient of cold pressor pain or the consoling partner undergoing EEG. EEG activity was recorded from the consoling partner during baseline and pain conditions (with and without consoling touch) to investigate how observation of a partner in pain could alter mu rhythms.

**Results:** Observing a romantic partner in pain significantly enhanced mu at electrodes C4 and Cz, yet addition of consoling touch did not significantly change mu rhythms. Personal Distress was positively associated with mu rhythm enhancement at electrode C4 during pain without consoling touch (*r* = .576, *p* = .008).

**Conclusion:** Viewing a romantic partner in pain significantly increased mu rhythms, that was not further changed by offering consoling touch. The association between increases in mu rhythms and Personal Distress in the current study suggests that self-oriented distress may be especially relevant to sensorimotor cortex responses when witnessing a partner in pain, particularly when consoling touch is not possible.

## Introduction

Observing a loved one in pain can evoke cognitive, emotional, and physiological responses that influence caregiving behaviors. Empathy is “understanding a person from their frame of reference rather than one’s own, or vicariously experiencing that person’s feelings, perceptions, and thoughts”[1]. This multifaceted construct is further categorized into two distinct forms: *trait empathy*, an enduring and intrinsic personal characteristic, and *state empathy*, denoting a situational and context-specific response to particular circumstances.

Romantic partners are among the most salient social relationships making them an important context for studying empathy and pain. Social touch influences our emotional state, offering relief from distress or discomfort across diverse contexts[8]. Prafull Jaltare, Sanfrinnin, and Torta demonstrated that handholding can modulate the spread of hypersensitivity robustly[12]. Observing another person’s pain can alter empathetic response and cortical activities. The level of empathy someone has can predict how much their primary somatosensory cortex activates when observing both non-painful and painful stimuli being applied to another person’s hand[8]. In a study using a fMRI, when shown pictures of their romantic partner, pain perception was attenuated and decreased blood oxygen level dependent activity in regions associated with pain processing was noted particularly in the left and right posterior insula [25]. These findings suggest that close interpersonal relationships influence the subjective pain experience.

A meta-analytic review of neuroimaging studies investigating empathy for pain indicated that empathy for pain shares neural substrates with pain processing[15]. This includes bilateral anterior insula and medial/anterior ACC processing [15]. A separate meta-analysis of 128 neuroimaging studies indicated that the AI, MCC, postcentral gyrus, thalamus, amygdala, brainstem, and inferior parietal lobe were all implicated in the cortical processing of empathy[22]. These meta-analytic insights paint the picture that empathy is diffusely processed across the cortex. Electroencephalography (EEG) presents an opportunity to examine cortical responses to empathy for a loved one in pain with high temporal resolution. One EEG-based cortical marker for pain and empathy is the mu rhythm. Initially characterized by Gastaut and colleagues in 1954, the mu rhythm manifests as an 8–12 Hz wave observed symmetrically across the sensorimotor cortex. Attenuation in the alpha rhythm, especially over central cortical areas (also called mu suppression), correlates with activity in areas such as the inferior parietal lobule, the dorsal premotor cortex, and the somatosensory cortex[10,19]. Mu rhythm suppression is commonly interpreted as an electrophysiological neural marker of sensorimotor mirroring processes that have been implicated when we observe the actions of others, including observation of pain[2,20].

Although previous research has demonstrated a relationship between empathy and mu suppression, other literature indicates a relationship between mu enhancement and empathy. For example, Jelsone-Swain, McCommon, Turk, and Roach (2023) found mixed findings with mu enhancement rather than suppression when painful images were presented[14]. Further, mu power during presentation of painful images was positively correlated with empathy scores. DiGirolamo and colleagues found no evidence to link mu suppression and trait empathy in a study of 252 participants[4]. The objective of the current study was to assess the relationship of mu activities to multidimensional empathy measures when participants are viewing a partner in pain.

## Methods

### Ethical Approvals and Recruitment

All study procedures were approved by the Tarleton State University Institutional Review Board (IRB). Participants provided informed consent prior to participation. All data were recorded at the same site at Tarleton State University. Participants were recruited using flyers on and off campus, a university email messaging system, and word of mouth. G*Power was used to compute a power analysis for a Pearson correlation with an alpha criterion of .05 and 80% power[6]. With a sample size of 20, this sample is powered to detect large correlations (r > .58). The total sample consisted of 52 participants (26 romantic couples). Four couples were excluded due to noisy EEG data, and two couples were excluded due to missing a final MPQ-SF score for one participant. This left a remaining total of 40 participants in the study (20 couples).

### Exclusion and Inclusion Criteria

Heterosexual couples were recruited if both individuals were aged 18 years or older and exhibited right-hand dominance. Participants were excluded if they (a) had a pacemaker, (b) were currently pregnant or possibly pregnant, (c) had a chronic pain condition, (d) engaged in athletic activities, were members of ROTC, or served in the military, (e) had experienced a concussion within the past two years, or (f) had received diagnoses of neurological disorder or psychological conditions.

### Measures

#### Edinburgh Handedness Inventory-Short Form

The Edinburgh Handedness Inventory (EHI) assessed handedness[24]. The EHI-Short Form consists of 4 items that ask individuals to indicate their preference for using either the right or left hand in specific activities. Scores on this measure can range from −100 to +100, with higher positive scores indicating stronger right-hand preference. Scores between +60 and +100 are generally classified as right-handed.

#### Short Form McGill Pain Questionnaire (SF-MPQ)

The SF-MPQ, comprised of 17 questions, has been utilized to assess various aspects of pain, including its intensity, quantity, and quality [16]The first 11 questions capture sensory descriptors of pain, such as throbbing, shooting, and stabbing, with ratings ranging from 0- no pain, 1- mild, 2- moderate, 3-severe[16]. Questions 12-15 delve into emotional/affective descriptors of pain, like tiring/exhausting and fearful, also rated on the same scale [16]. Participants gauge their pain intensity using a visual analogue scale, marking their pain level on a continuum from “no pain” to “worst possible pain” [16]. Finally, the Present Pain Intensity (PPI) index prompts participants to rate their current pain intensity from “0- no pain” to “5-excruciating” [16]. Data from the SF-MPQ were aggregated to yield scores for sensory pain (items 1-11), emotional pain (items 12-15), visual analog scale (question 16), and PPI (question 17). Scores of 0 represent no pain, while higher scores represent more pain.

#### Toronto Empathy Questionnaire (TEQ)

Developed by Davis and Goldberg, The Toronto Empathy Questionnaire (TEQ) is a psychological assessment tool designed to measure empathy in individuals[21]. The questionnaire consists of 16 scenarios, each describing a situation where someone requires help or support. Respondents rate their level of empathic concern for each scenario on a scale ranging from 0-never (low empathy) to 4-always (high empathy) [21]. Negatively worded items are reverse-coded. Items are summed for each participant. High scores on the TEQ suggest strong empathy-related abilities or tendencies. In the current study, the TEQ demonstrated good internal consistency across items (Cronbach’s α = .869, SE = .036, 95% CI [.799, .939].

#### Interpersonal Reactivity Index (IRI)

The Interpersonal Reactivity Index (IRI) is a 28-item multidimensional self-report questionnaire that measures trait empathy[3] Instead of viewing empathy as a singular, one-dimensional concept, the IRI assesses empathy as a collection of related but distinct dimensions. The IRI consists of four subscales (7 questions each), each measuring a different component of empathy: Perspective Taking, Fantasy, Empathic Concern, and Personal Distress. For each question, the participant must rate on a five point Likert scale of A - “Does Not Describe Me Well” to E - “Describes Me Very Well.” Select items are reversed scored. Scores on each 7-item subscale are then calculated by summing the participant’s responses to the respective items, with higher scores indicating greater levels of the measured empathy dimension. Internal consistency was evaluated for each IRI subscale using Cronbach’s alpha: For the subscale Fantasy, Cronbach’s α = .766, SE = .082, 95% CI [.606, .926], for the subscale Empathic Concern, Cronbach’s α = .806, SE = .056, 95% CI [.696, .916], for the subscale Perspective-Taking, Cronbach’s α = .787, SE = .061, 95% CI [.667, .906] and for the subscale Personal Distress, (Cronbach’s α = .642, SE = .134, 95% CI [.379, .906].

#### Cold Pressor Task

For the current study, a cold water therapy wrap system (Aqua Relief System) was re-designed for use as a cold pressor apparatus to incite acute pain. It consists of the base compartment, which holds the water and ice, from which stems a hose with an arm cuff. The ice water circulates in the cuff to ensure a constant cold temperature. Water was cooled with an aquarium chiller (SDLOOL) and nugget ice (Humhold Ice Maker). Each recording was 5 minutes long with a 15-minute break after each cold pressor administration and a water temperature range close to 1 degree Celsius. Previous research has indicated that this temperature produces the highest pain scores compared to other temperatures[5,17].

#### Electroencephalogram Recording

EEG activity was recorded in a closed sound attenuating chamber with LED lights using NetStation acquisition software and a Magstim EGI GES 400 amplifier. HydroCel Geodesic 256 electrode sensor nets were prepared in 2 liters of water with Potassium Chloride and Johnson’s Baby Shampoo. Electrode nets were placed using the international 10-20 system and referenced to all electrodes. Data were recorded at a sampling rate of 1096 Hertz. Following application of the sensor net, electrode impedance was checked and maintained below 50 kΩ before and during recording.

### Procedure

Upon arrival at the lab, participants completed a comprehensive survey with (a) informed consent and exclusion criteria, (b) demographic questions regarding participants’ age (continuous), biological sex (male, female, intersex), gender identity (cisgender woman, cisgender male, transgender woman, transgender man, nonbinary, gender non-conforming), race (White, Black or African American, American Indian or Native American or Alaska Native, Asian/Asian American, Native Hawaiian, Pacific Islander, Hispanic or Latino, Middle Eastern or North African, Other), ethnicity (Hispanic or Latino, not Hispanic or Latino), marital status (single-never married, widow, widower, single-divorced, not married but cohabitating, married but not cohabitating, married and cohabitating), and how long the couple has been in a relationship (continuous) (c) the Edinburgh Handedness Inventory, (d) Toronto Empathy Questionnaire, (e) Interpersonal Reactivity Index, and (f) Short Form McGill Pain Questionnaire (SF-MPQ).

The experimenter explained that either the male or female (randomized) would be subject to the pain stimulus, while the other would offer a consoling touch while wearing an electroencephalogram (EEG) cap. Then, the designated individual (male or female-counterbalanced) was fitted with the EEG net, preceded by an impedance check to ensure optimal data quality. After this preparatory phase, the EEG recording began, encompassing four recordings: (1) baseline with no pain and no consoling touch (5 minutes), (2) baseline with no pain and consoling touch (5 minutes), (3) in pain with no consoling touch (5 minutes), and (4) in pain with consoling touch (5 minutes). The order of the 5-minute recordings differed based on two groups, with one group linearly progressing through the four recordings and the other group progressing through baseline with no pain and consoling touch, baseline with no pain and no consoling touch, in pain with consoling touch, then in pain with no consoling touch. After each cold pressor recording, there was a break of 15 minutes. Following the final recording, participants completed the MPQ-SF again, and the EEG cap was removed, and participants exited the laboratory (see Figure 1).

**Figure 1.**
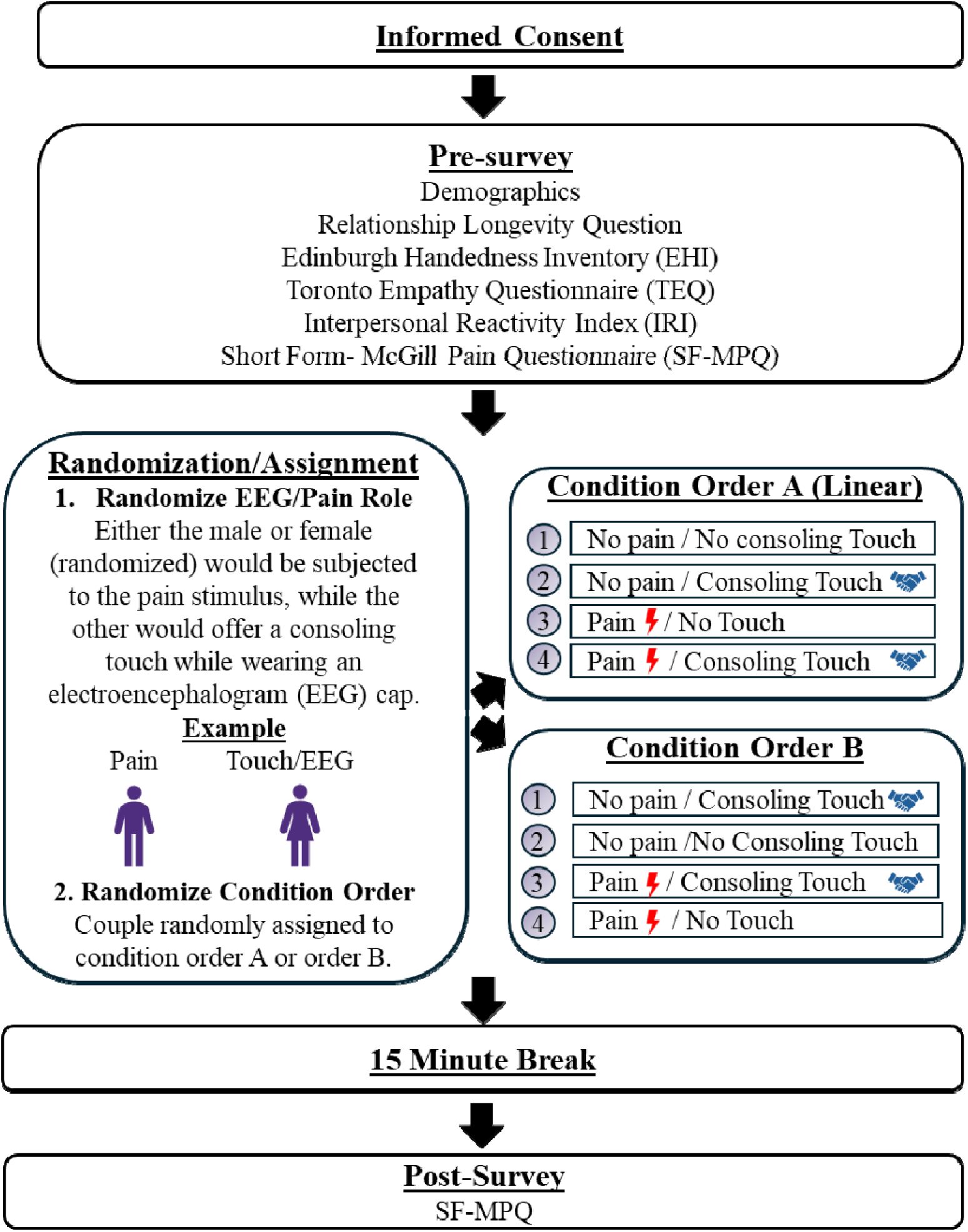
Experimental Design Flowchart. Participants completed informed consent and demographic questions. In addition to a question about relationship longevity, participants completed baseline questionnaires (EHI, TEQ, IRI, and SF-MPQ). Members of each of the 20 couples were randomized to either be (a) the recipient of a an acute pain stimulus (cold pressor) or (b) the observer who offered consoling touch and underwent an EEG recording. Couples were randomized to one of two experimental condition orders. EEG was recorded for 5 minutes during each of the experimental conditions. A 15-minute break was allotted after each cold pressor administration. To conclude, the SF-MPQ was administered again.

#### EEG Processing and Mu Calculations

Using NetStation software, the raw EEG sensor data first underwent pre-processing where the data were filtered with a bandpass filter (.10-50 Hz) and notch filter (60 Hz), data were segmented (3000 ms time bins), and artifacts were detected. Artifacts were rejected (eye blinks, bad channels, and eye movements) and data were reviewed in segmented time bins to check for artifacts such as edge artifacts. Bad channels were replaced, data were averaged, and baseline correction was applied. Channels were marked bad for all segments if they were bad for greater than 20% of segments and contained more than 10 bad channels, contained an eye blink, or eye movements. Data from the alpha frequency range (8-13Hz) were extracted for analysis.

To quantify mu rhythm activity, a baseline measure of EEG power at central sites for comparison to the EEG power during the experimental conditions of interest is needed (Hobson & Bishop, 2016). Mu suppression or enhancement can be assessed by comparing the power during experimental conditions to that in the baseline condition. This ratio approach accounts for variations in absolute mu power due to factors like scalp thickness and electrode impedance. A log transformation is applied for analysis as follows:

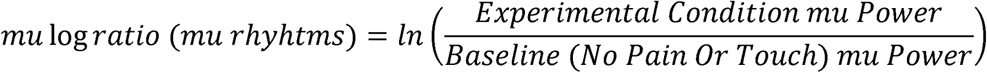

A log ratio below zero signifies mu suppression, a score of 0 represents no change relative to baseline, and values above zero suggest an increase in mu power/enhancement of mu when compared to baseline[18,19]. Figures were made using Pandion Plots[23].

## Results

### Participant Characteristics

Data were analyzed in JASP [13]and are presented as mean ± SD for continuous variables and frequency and percent of sample for categorical variables. Participant demographics are depicted in Table 1. Relationship longevity ranged from 4 months to 24 years. The average relationship length was 44.45 months or approximately 3.7 years (*SD* = 77.28; *Range* = 4-288 months).

**TABLE 1.** Participant Characteristics.

| Continuous Variable | Total Sample |  | EEG Group |  |
| --- | --- | --- | --- | --- |
|  | Mean | SD | Mean | SD |
| Age | 25.51 | 12.04 | 24.25 | 10.49 |
| Categorical Variables | Total Sample |  | EEG Group |  |
|  | N | % of Sample | N | % of Group |
| **Biological Sex** |  |  |  |  |
| Male | 20 | 50% | 10 | 50% |
| Female | 20 | 50% | 10 | 50% |
| Intersex | 0 | 0% | 0 | 0% |
| **Gender** |  |  |  |  |
| Cisgender Woman | 20 | 50% | 10 | 50% |
| Cisgender Man | 19 | 48% | 9 | 45% |
| Transgender Woman | 0 | 0% | 0 | 0% |
| Transgender Man | 0 | 0% | 0 | 0% |
| Nonbinary | 1 | 3% | 1 | 5% |
| Gender Non-Conforming | 0 | 0% | 0 | 0% |
| **Race** |  |  |  |  |
| White | 34 | 85% | 16 | 80% |
| Black or African American | 0 | 0% | 0 | 0% |
| Asian/Asian American | 0 | 0% | 0 | 0% |
| Native Hawaiian | 0 | 0% | 0 | 0% |
| Pacific Islander | 0 | 0% | 0 | 0% |
| Hispanic or Latino | 6 | 15% | 4 | 20% |
| Middle Eastern or North African | 0 | 0% | 0 | 0% |
| Other | 0 | 0% | 0 | 0% |
| **Ethnicity** |  |  |  |  |
| Hispanic or Latino | 6 | 15% | 5 | 25% |
| Not Hispanic or Latino | 34 | 85% | 15 | 75% |
| **Marital Status** |  |  |  |  |
| Never Married | 29 | 73% | 14 | 70% |
| Widow | 0 | 0% | 0 | 0% |
| Widower | 0 | 0% | 0 | 0% |
| Single-Divorced | 0 | 0% | 0 | 0% |
| Not Married but Cohabiting | 7 | 18% | 4 | 20% |
| Married but not Cohabiting | 0 | 0% | 0 | 0% |
| Married and Cohabiting | 4 | 10% | 2 | 10% |

### Edinburgh Handedness Inventory Results

Laterality quotient scores ranged from 62.50 to 100.00, indicating that all EEG participants fell within the right-handed range (*M* = 94.38, *SD* = 12.48).

### Short Form McGill Pain Questionnaire Results

The SF-MPQ was administered to participants (*n* = 20) who received the acute pain stimulus at baseline and immediately following the completion of all EEG recordings (Table 2).

**Table 2.** SF-MPQ Descriptive Statistics.

|  | Sensory<br>Baseline | Sensory<br>Post | Affective<br>Baseline | Affective<br>Post | VAS<br>Baseline | VAS<br>Post | PPI<br>Baseline | PPI<br>Post |
| --- | --- | --- | --- | --- | --- | --- | --- | --- |
| Mean | 1.90 | 6.30 | 0.70 | 0.85 | 0.65 | 3.60 | 0.35 | 1.90 |
| SD | 4.59 | 4.98 | 2.03 | 1.73 | 1.50 | 2.42 | 0.81 | 1.07 |
| Minimum | 0.00 | 0.00 | 0.00 | 0.00 | 0.00 | 0.00 | 0.00 | 0.00 |
| Maximum | 19.00 | 18.00 | 9.00 | 7.00 | 6.00 | 8.00 | 3.00 | 4.00 |

Paired samples t-tests for each SF_MPQ construct computed to analyze potential change from baseline to cold pressor administration revealed that implementation of the cold pressor acute pain stimulus significantly increased the sensory dimension scores, VAS scores, and PPI scores, *p* < .05 (Table 3).

**Table 3.** SF-MPQ T-Tests.

|  |  |  | *t* | *df* | *p* | Cohen's *d* | 95% CI for |  |
| --- | --- | --- | --- | --- | --- | --- | --- | --- |
|  |  |  |  |  |  |  | Lower | Upper |
| Sensory Baseline | - | Sensory Post | -2.664 | 19 | *0.015 | -0.596 | -1.066 | -0.112 |
| Affective Baseline | - | Affective Post | -0.234 | 19 | 0.818 | -0.052 | -0.49 | 0.387 |
| VAS Baseline | - | VAS Post | -3.904 | 19 | *< .001 | -0.873 | -1.383 | -0.347 |
| PPI Baseline | - | PPI Post | -4.233 | 19 | *< .001 | -0.947 | -1.468 | -0.408 |

\*Significantly different at alpha criterion of .05.

Mixed ANOVAs were computed to determine if there were biological sex differences on MPQ scores. MPQ scores in each domain were run as the repeated measures factor while sex was run as the between subjects factor. For SF-MPQ scores over time, there was no main effect of sex on sensory scores [*F* (1, 18) = 1.61, *p* =.220, η*p²* = .082], on affective scores [*F* (1, 18) = 1.46, *p* = .243, η*p²* = .075], or on VAS [*F* (1, 18) = 1.30, *p* =.269, η*p²* = .067]. However, there was a main effect of sex on PPI with males reporting significantly higher PPI scores than females [*F* (1, 18) = 5.32, *p* =.033, η*p²* = .228].

### Toronto Empathy Questionnaire Results

Trait empathy was assessed using the Toronto Empathy Questionnaire (TEQ). Total TEQ scores for participants receiving the EEG (*n* = 20) ranged from 36.00 to 59.00 (possible range of 0 - 64), with higher scores indicating greater levels of trait empathy. The mean TEQ score was 48.10 (*SD* = 7.39), suggesting moderate to high levels of self-reported trait empathy among participants. An ANOVA computed to assess potential sex differences in TEQ scores revealed that males and females reported significantly different empathy on the TEQ, *F* (1,18) = 14.38, *p* = .001, η*p²* = .444. Specifically, females reported higher empathy on the TEQ (*M* = 52.90, *SD* = 6.12) than males (*M* = 43.30, *SD* = 5.17).

### Interpersonal Reactivity Index Results

Subscale scores were calculated for the four standard IRI components (with a possible mean of up to 28; See Fig. 2): Perspective-Taking, Empathic Concern, Fantasy, and Personal Distress. Perspective - Taking scores ranged from 14.00 to 27.00 (*M* = 19.55, *SD* = 4.27). Fantasy scores ranged from 5.00 to 25.00 (*M* = 17.00, *SD* = 5.31). Empathic Concern scores ranged from 10.00 to 28.00 (*M* = 20.20, *SD* = 5.09). Personal Distress scores ranged from 1.00 to 18.00 (*M* = 10.35, *SD* = 4.53).

**Figure 2.**
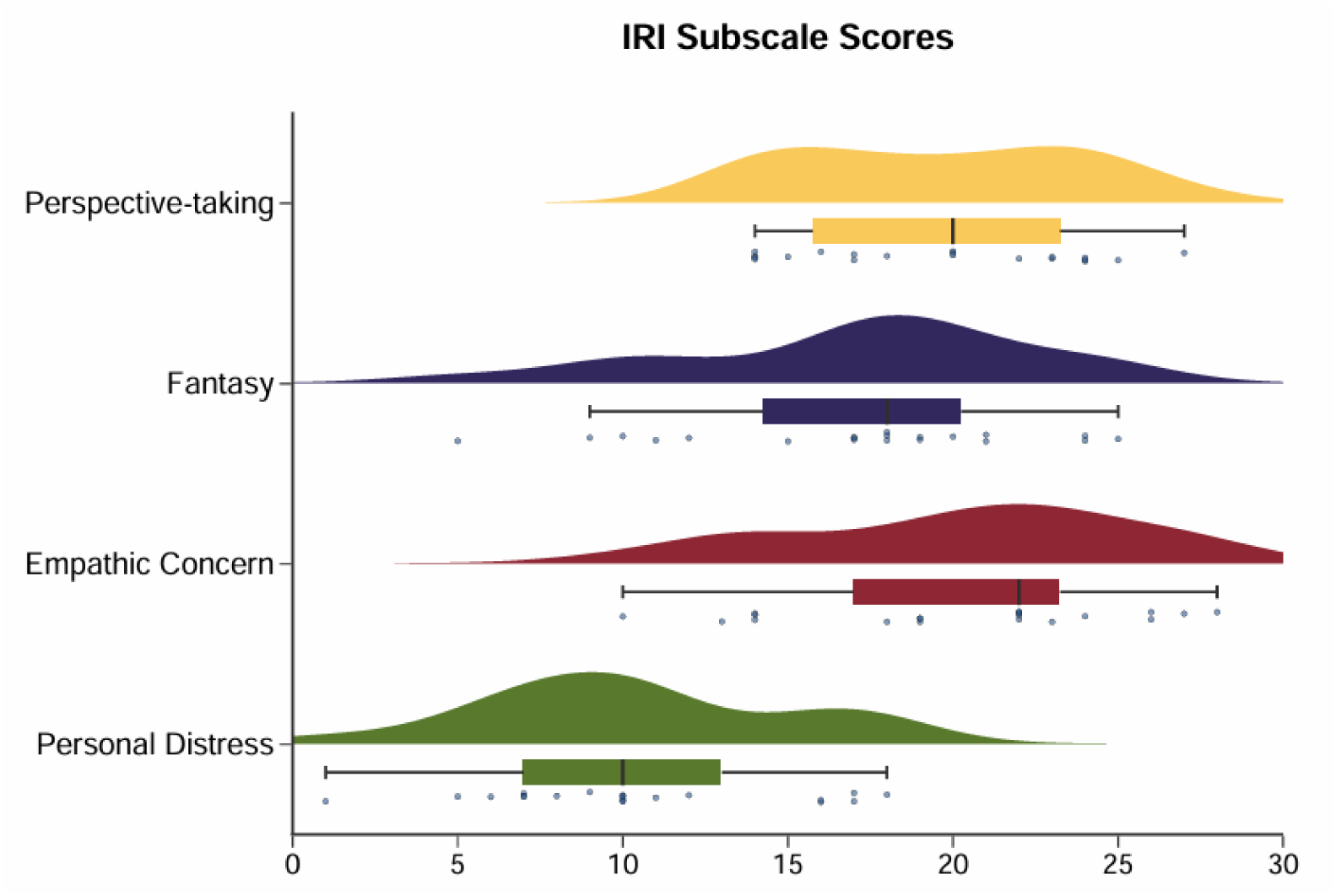
Raincloud Plot of IRI subscales for the EEG group. Mean scores (+-SD), spread, and data points are indicated for each of the four subscales of the IRI. Higher scores reflect higher levels of the empathy dimension.

An ANOVA computed to assess potential sex differences in IRI scores revealed that males and females reported significantly different empathy on the IRI overall, *F* (1,18) = 26.91, *p* < .001, η*p²* = .599. Specifically, females reported higher empathy (*M* = 77.60, *SD* = 8.02) than males (*M* = 56.60, *SD* = 9.98). For each of the four subscales, females reported higher empathy than males, *p* <.05.

### EEG Results

To assess neural responses associated with empathy for pain, the mu rhythm was quantified for each experimental condition relative to baseline (no pain, no touch) at the central electrodes of interest and log transformed (See Table 4).

**Table 4.**
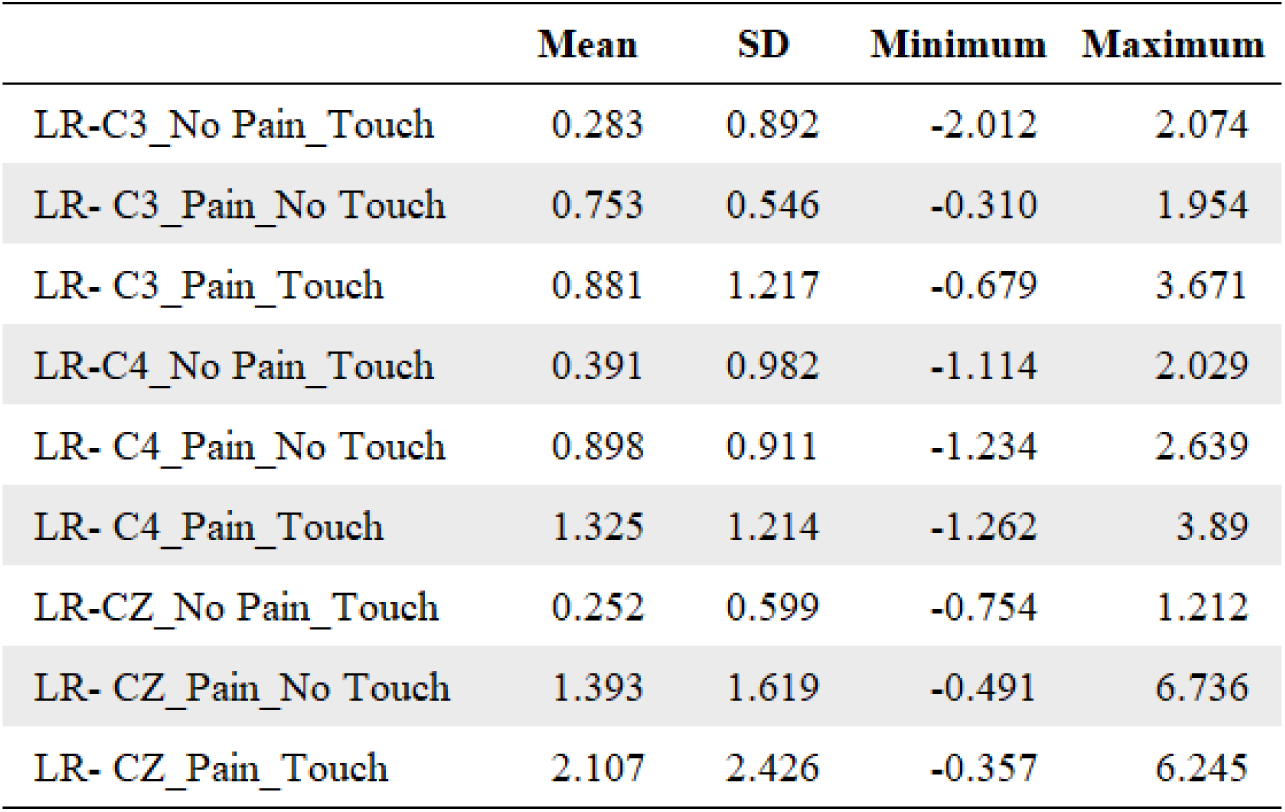
EEG Mu Rhythm Descriptives.

Repeated measures ANOVAs were computed for each electrode of interest to assess how mu rhythms changed with varying input to the nervous system (pain and touch). For C3, there was no significant effect of condition, *F* (2, 38) = 3.97, *p* = .065, ηp² = .134. In contrast, there was a significant effect of condition for C4, *F* (2, 38) = 8.74, *p* = .002, η*p²* = .286. Specifically, pain/no touch (*M* = .90, *SD* = .91) and pain/touch (*M* = 1.33, *SD* = 1.21) were significantly higher than no pain/touch (*M* = .39, *SD* = .98), indicating mu enhancement. There was also a significant main effect of condition for Cz, *F* (2, 38) = 17.50, *p* < .001, η*p²* = .324. Again, pain/no touch (*M* = 1.39, *SD* = 1.62) and pain/touch (*M* = 2.11, *SD* = 2.43) were significantly higher than no pain/touch (*M* = .25, *SD* = .60), indicating mu enhancement.

To analyze any potential sex differences in mu rhythms, mixed ANOVAs were run with sex as the between subjects variable and mu scores across conditions as the within subjects variables. There were no sex differences in mu over time for electrode C3 [*F* (1, 18) = 1.33, *p* =. 263, η*p²* = .069], or for electrode C4 [*F* (1, 18) = 5.40, *p* = .112, η*p²* = .134]. There was a significant sex difference for electrode Cz [*F* (1, 18) = 9.65, *p* = .006, η*p²* = .349].

#### Mu Rhythms and Empathy

Pearson’s correlations were computed to assess the relationship between mu scores (relative to baseline) in the electrodes of interest and empathy as measured by the TEQ. To account for multiple comparisons, an alpha criterion correction was computed by dividing the original alpha criterion of .05 by the number of TEQ comparisons (2 pain conditions X 3 electrodes), resulting in an adjusted α = .008. During pain and consoling touch, TEQ was positively correlated with mu scores (relative to baseline) in electrode Cz, *r* = .540, *p* = .014, however, this association did not meet the adjusted threshold for statistical significance (see Table 5 and Figure 3). Because positive log-ratio values of mu indicate enhancement of mu compared to baseline, these values indicate mu enhancement rather than mu suppression.

**Figure 3.**
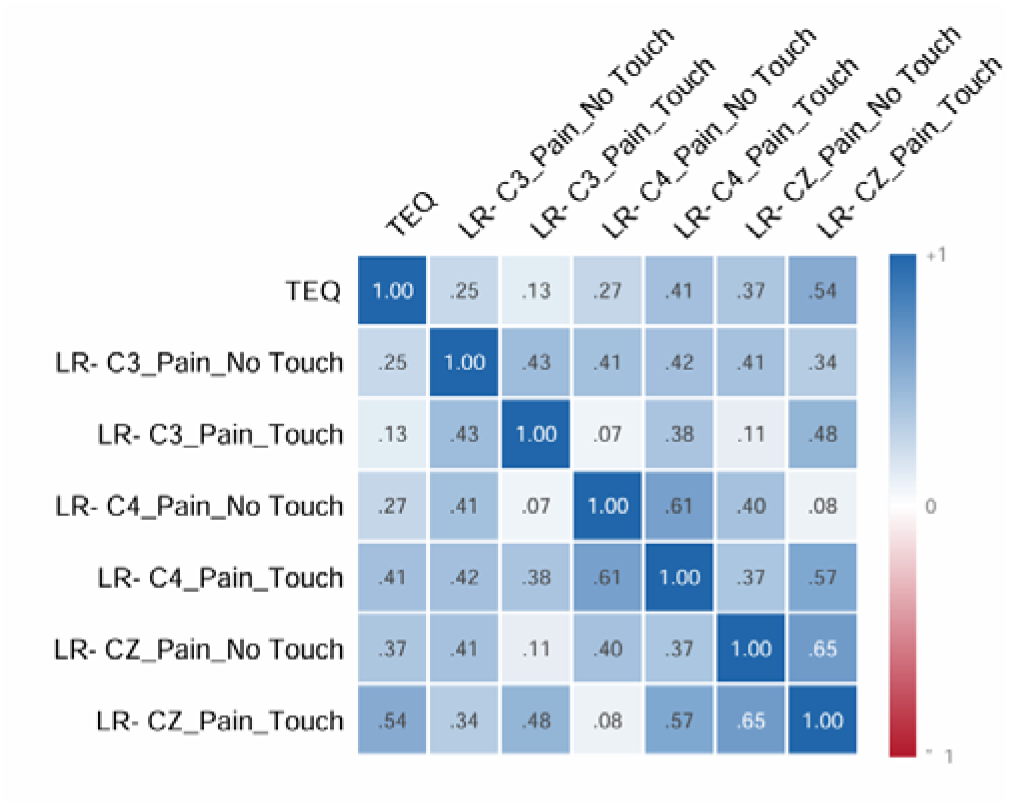
Heat Map of Correlations between TEQ and Mu Rhythms. A heat map of correlations between the Toronto Empathy Questionnaire and each mu rhythm (C3, Cz, C4) during pain/no touch and pain/touch. The intensity of the colors represents the magnitude of the correlation.

**Table 5.** Pearson’s Correlations-TEQ and Mu Rhythms.

| **Mu Rhythms Compared to TEQ** | **Pearson's r** | **p** |
| --- | --- | --- |
| LR- C3_Pain_No Touch | 0.248 | 0.291 |
| LR- C3_Pain_Touch | 0.127 | 0.595 |
| LR- CZ_Pain_No Touch | 0.366 | 0.112 |
| LR- CZ_Pain_Touch | 0.540 | **0.014\*** |
| LR- C4_Pain_No Touch | 0.265 | 0.258 |
| LR- C4_Pain_Touch | 0.414 | 0.070 |

\*Significant correlation at $\alpha = .05$ , but not adjusted $\alpha = .008$

Pearson’s correlations were computed to assess the potential relationship between mu rhythms (relative to baseline) in the electrodes of interest and empathy as measured by the IRI subscales. Each IRI subscale was evaluated separately, with an adjusted alpha criterion of .008 (.05/6). At electrode Cz during pain and consoling touch there was a positive correlation with IRI on the empathic concern subscale, *r* = .538, *p* = .014, however, this association did not meet the adjusted threshold for statistical significance. During the pain without consoling touch condition at electrode C4, there was a significant positive correlation with Personal Distress, *r* = .576, *p* = .008 (See Table 6). These positive log-ratio values of mu indicate enhancement of mu compared to baseline.

**Table 6.** Pearson’s Correlations-IRI Subscales and Mu Rhythms.

| Mu Rhythms | Perspective Taking |  | Fantasy |  | Empathic Concern |  | Personal Distress |  |
| --- | --- | --- | --- | --- | --- | --- | --- | --- |
|  | *r* | *p* | *r* | *p* | *r* | *p* | *r* | *p* |
| LR- C3_Pain_No Touch | 0.025 | 0.915 | 0.080 | 0.738 | 0.320 | 0.169 | 0.075 | 0.753 |
| LR- C3_Pain_Touch | -0.141 | 0.553 | -0.084 | 0.725 | 0.143 | 0.547 | -0.181 | 0.445 |
| LR- C4_Pain_No Touch | 0.023 | 0.922 | 0.167 | 0.483 | 0.287 | 0.220 | 0.576 | **0.008\*\*** |
| LR- C4_Pain_Touch | 0.123 | 0.607 | -0.005 | 0.985 | 0.382 | 0.096 | 0.271 | 0.248 |
| LR- CZ_Pain_No Touch | 0.150 | 0.527 | 0.151 | 0.525 | 0.347 | 0.134 | 0.362 | 0.117 |
| LR- CZ_Pain_Touch | 0.210 | 0.375 | 0.140 | 0.557 | 0.538 | **0.014\*** | 0.083 | 0.728 |

\*Significant correlation at $\alpha = .05$
\*\*Significant at adjusted $\alpha = .008$

**Figure 4.**
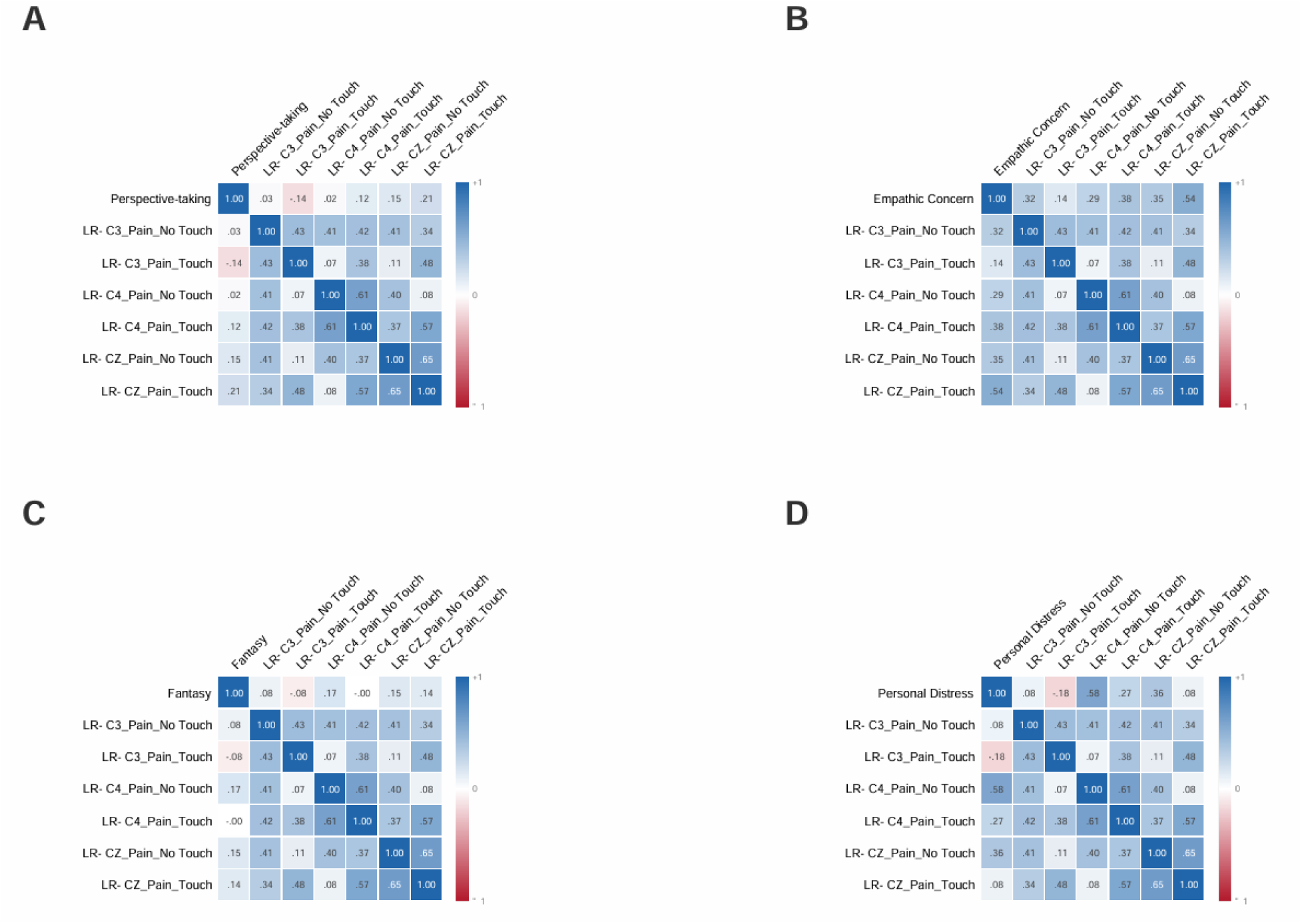
Heat Maps of Correlations between IRI subscales and Mu Rhythms. Heat maps of correlations between the Interpersonal Reactivity Index Subscales (A-Perspective Taking, B-Empathic Concern, C-Fantasy, and D-Personal Distress) and each mu rhythm (C3, Cz, C4) during pain/no touch and pain/touch. The intensity of the colors represents the magnitude of the correlation.

## Discussion

The present study examined whether multidimensional trait empathy was associated with mu rhythms while participants observed their romantic partner experiencing experimentally induced pain, with and without consoling touch. Participants reported moderate to high levels of empathy in response to the scenarios provided by the questionnaires, indicating a high disposition in the sample towards empathetic responding. Consistent with previous studies, females reported significantly higher empathy in every empathy domain in the current study.

Previous research has shown that observing another individual experiencing pain can elicit increased mu suppression, suggesting greater empathic engagement and sensorimotor simulation of another person’s experience[11]. In the current study, overall, observation of a partner in pain (but not addition of consoling touch) enhanced mu. At the C4 and Cz (midline) electrodes, mu rhythms significantly increased with the observation of the application of a pain stimulus to the partner, but not with the addition of consoling touch to the partner. In other words, the addition of consoling touch alone did not produce any significant changes in mu rhythms. The observation of the partner’s pain experience (rather than consoling touch), increased mu. Although females reported greater empathy than males, sex differences in mu were limited to the Cz electrode.

Prior research has indicated a mixed relationship between mu rhythms and empathy. In the current study at electrode Cz, greater TEQ and Empathic Concern scores were associated with greater mu during the pain/touch condition, however, neither association remained statistically significant after correction for multiple comparisons. In contrast, greater Personal Distress was associated with greater C4 mu during the pain/no touch condition and was significant after alpha correction. This distinction may be important, because personal distress is a self-oriented response to the suffering of the partner in contrast with empathic concern which reflects other-oriented attention. Individuals who experience greater personal distress when witnessing another’s pain may exhibit greater activity in the sensorimotor cortex when observing a romantic partner in pain, especially when not able to provide physical comfort. Although consoling touch did not independently change mu, the pattern of associations differed between the touch and no touch conditions.

Existing literature indicated that physical support from a romantic partner can attenuate the pain experience and alter the neural response. Goldstein et al. (2018), for example, demonstrated that consoling touch during pain increased brain-to-brain coupling and greater coupling weas associated with analgesia and empathetic accuracy[9]. Much of the literature in this domain has focused on the individual receiving pain rather than the partner observing the pain. The present findings provide a complementary perspective by focusing on the individual observing the pain. This relationship is dependent on the dimension of empathy rather than empathy overall and is dependent on the context in which the partner’s pain is observed. A unique aspect of the present study is the focus on the observation of a partner in pain, and the focus on the multidimensional subscale measures of empathy which may be differentially contributing to the mu response.

Ferguson et al. used EEG to investigate whether we use the same core empathy networks for physical versus social pain and how they change from adolescence into adulthood[7]. The authors found that mu rhythm suppression increased when participants viewed images of others in pain, both socially and physically[7]The current study extended this paradigm beyond pictures. We found that viewing a partner in pain in real time increased mu rhythms, and significantly as it pertained to personal distress. The current findings also differ from those presented by Peled-Avron et al. who investigated mu activity during touch in romantic partners receiving heat pain stimulation and the relationship with situational empathy in a similar sample size (18 couples) [19]. They reported a finding of greater mu suppression in C3, C4, and Cz during consoling touch, and C4 suppression was related to a construct of situational empathy. Previous findings and the current findings support the idea that neural activity related to empathy may be dimension and context dependent. Future studies are needed to clarify the role of mu suppression or enhancement before mu rhythms are utilized in a clinical or diagnostic setting.

## Limitations and Future Research

The sample consisted exclusively of heterosexual romantic couples, which may limit generalizability of the findings to other relationships. The demographic composition of the sample was also relatively homogenous. The current sample was powered for moderate to large correlational results with a sample size slightly larger than other similar published studies [19,25]. Small correlations may have been undetectable.

## Acknowledgements

The authors wish to thank Honors thesis committee member Dr. Eileen Faulkenberry for her valuable contributions to the conceptualization of the design for this project. We are grateful to the study participants for their time participating in the study-without whom this work would not be possible.

## Data Availability Statement

Data are available at Open Science Framework: https://osf.io/4qswj/overview?view_only=830a99a6f9f244d5ad87b7601c91d8a7

## Authorship Contributions

**Sarah Love:** Conceptualization, Methodology, Formal Analysis, Investigation, Writing-Original Draft, Visualization, Project Administration. **Lauren Rowden-** Formal Analysis, Investigation, Writing-Review & Editing. **Bryten Ballenger-** Formal Analysis, Investigation, Writing-Review & Editing. **Amber Harris Bozer-** Conceptualization, Methodology, Software, Formal Analysis, Investigation, Resources, Writing- Original Draft, Writing- Review & Editing, Visualization, Supervision, Project Administration, Funding Acquisition.

